# Laterality defect of the heart in non-teleost fish

**DOI:** 10.1101/2023.06.13.544822

**Authors:** Myrte M. Huijskes, José M. Icardo, Bram F. Coolen, Bjarke Jensen

## Abstract

Dextrocardia is a rare congenital malformation in humans in which most of the heart mass is positioned in the right hemithorax rather than on the left. The heart itself may be normal and dextrocardia is sometimes diagnosed during non-related explorations. A few reports have documented atypical positions of the cardiac chambers in farmed teleost fish. Here, we report the casual finding of a left-right mirrored heart ventricle in a 85 cm long spiny dogfish (*Squalus acanthias*) that was caught in the wild. Macroscopic observations showed an outflow tract originating from the left side of the ventricular mass, rather than from the right. Internal inspection revealed the normal structures, including a looped cavity on the inner curvature of which was a large trabeculation, or the bulboventricular fold. MRI data acquired at 0.7 mm isotropic resolution and subsequent 3D-modeling revealed the atrioventricular canal was to the right of the bulboventricular fold, rather than on the left. The atrium and sinus venosus had the normal midline position and gross shape. Spurred by finding of the left-right mirrored shark ventricle, we revisit our previously published material on farmed Adriatic sturgeon (*Acipenser naccarii*), a non-teleost bony fish. We found several alevins with inverted (L-loop) hearts, amounting to an approximate incidence of 1-2 %. Additionally, an adult sturgeon measuring 90 cm in length showed abnormal topology of the cardiac chambers, but normal position of the abdominal organs. In conclusion, left-right mirrored hearts, a setting that resembles human dextrocardia, can occur in both farmed and wild non-teleost fish.

## Introduction

While dissecting formalin-fixed spiny dogfish (*Squalus acanthias*) as part of an elective course for medical students at our institute, we came across one case of apparent situs inversus in a male specimen that was outwardly mostly normal. Due to the dissection and preparation of this specimen, several organs were displaced, but the heart was in situ when it was realized that the ventricle was left-right mirrored. A setting resembling situs inversus has been reported in inbred strains of platyfish (Baker‐Cohen, 1961) and farmed salmonids can exhibit various cardiac malformations, including abnormal chamber topology (Poppe et al 2002; Frisk et al 2020). Given the high degree of left-right symmetry of the heart in the vast majority of bony fish (Icardo 2017), even when a substantial conus arteriosus is present (Icardo 2006), it is challenging to assess whether the cardiac chambers may be affected by a laterality defect (Baker‐Cohen, 1961). Early in development, however, bony fish hearts exhibit much more asymmetry than in adult stages, both anatomically and molecularly (Icardo et al 2004; Noël et al 2013; Smith and Uribe 2021). In humans, dextrocardia together with situs inversus of other organs can be a casual finding and only in about a quarter of the cases present with overt cardiac malformations (Bohun et al 2007). To our knowledge, this is the first report of a left-right mirrored heart in a cartilaginous fish. In addition, we revisited the material of a series of studies on the heart of the Adriatic sturgeon (Icardo et al 2002, 2004, 2009) and report left-right mirrored hearts in several alevins and in a specimen of approximately 90 cm total length (TL).

## Materials and methods

### Spiny dogfish

Six spiny dogfish of approximately 85 cm length and both sexes, previously fixed, were dissected as part of an elective course on vertebrate anatomy at Amsterdam UMC. The case specimen and an ostensibly normal shark were kept afterwards in a preservation fluid containing low concentrations of ethanol, glycerol, and phenol. For MRI, to wash out compounds in the preservation fluid that diminishes the tissue-free fluid contrast, the two sharks were placed under running tap water for three days. After that, over the weekend, they were stored at 5 °C in a sealed plastic bag before being MRI scanned on Whit Monday. They were scanned on a 3T Ingenia clinical MRI scanner (Philips, Best, the Netherlands) using a standard 16 channel anterior coil in combination with 12 posterior coil elements fixed in the MRI table. A high-resolution isotropic 3D T1-weighted gradient-echo sequence was used in order to suppress the fluid for optimal contrast. Specific sequence parameters were: TR/TE = 10/2.1 ms, flip angle = 20 degrees, FOV = 400 x 280 x 150 mm, resolution = 0.7 × 0.7 × 0.7 mm^3^, total scan time = 14 min. The MRI image stack of the case specimen is publicly available on MorphoSource (to come). We imported the image stack to the 3D software Amira (version 3D 2021.2, FEI SAS, Thermo Fisher Scientific). A Volume Rendering was made of the whole scan and a plane of sectioning was made using the Slice module. Next, labelling of the cardiac chambers and the atrioventricular canal was done in the Segmentation Editor module. From the label file, using the Generate Surface function, a surface file was made that was visualized using the Surface View function. This was done in the same window as the volume rendering whereby the heart model was projected into it.

### Adriatic sturgeons

Cardiac malformations in alevins of the autochthonous Adriatic sturgeon *Acipenser naccarii* (*A. naccarii* Bonaparte 1836) were found among specimens from previous studies (Icardo et al 2004, 2009). Briefly, alevins obtained from Sierra Nevada Fishery at Riofrío, Granada, Spain, were collected between days 1 and 28 post-hatching (dph). For scanning electron microscopy (SEM), the specimens were fixed in 3% glutaraldehyde in phosphate-buffered saline (PBS) for 3-5hr. Then, the samples were microdissected under a stereomicroscope, dehydrated in graded acetone, dried by the critical-point method with CO_2_ as the transitional fluid and coated with gold following routine procedures. Samples were observed with an Inspect S microscope (FEI Company) working at 15-20 KV. In addition, an unpublished case of laterality defect in a sturgeon of approximately 90 cm TL was found among the material of a previous study (Icardo et al 2002).

## Results and Discussion

The dissected sharks measured 85 cm TL. The stomach of the case specimen was full with partially digested fish and the testis were enlarged indicating that the shark had entered the reproductive stage (Figure 1A). There was bilateral agenesis of the spiracles. Most of the spiral intestine was prolapsed out of the cloaca. It was not clear whether the prolapse had already occurred when the shark was still alive. The liver was relatively large, yellowish in color, and comprised two large lobes but it lacked the third, smaller, right-sided medial lobe found in the other specimens. On the left cranial-medial side of the right liver lobe, a small fibrotic sack-like structure was found, which could be a remnant of the gall bladder. The stomach was right-sided instead of showing the normal left configuration. The pylorus curved to the left. From the pylorus, the small intestine curved cranially, then crossed to the left side and, finally, curved caudally. Both the spleen and pancreas had an abnormal morphology. Instead of the normal triangular shape, the spleen had an elongated shape, was located on the ventro-lateral side of the stomach and continued over the additional looping of the intestines. The pancreas followed a similar pattern, it was excessively elongate and followed the looping of the small intestine. The two pancreatic lobes were situated on the left side of the body. In aggregate, the visceral organs gave multiple indications of pronounced deviations from the normal topology (the MRI image stack can be viewed on MorphoSource).

**Figure 1.**
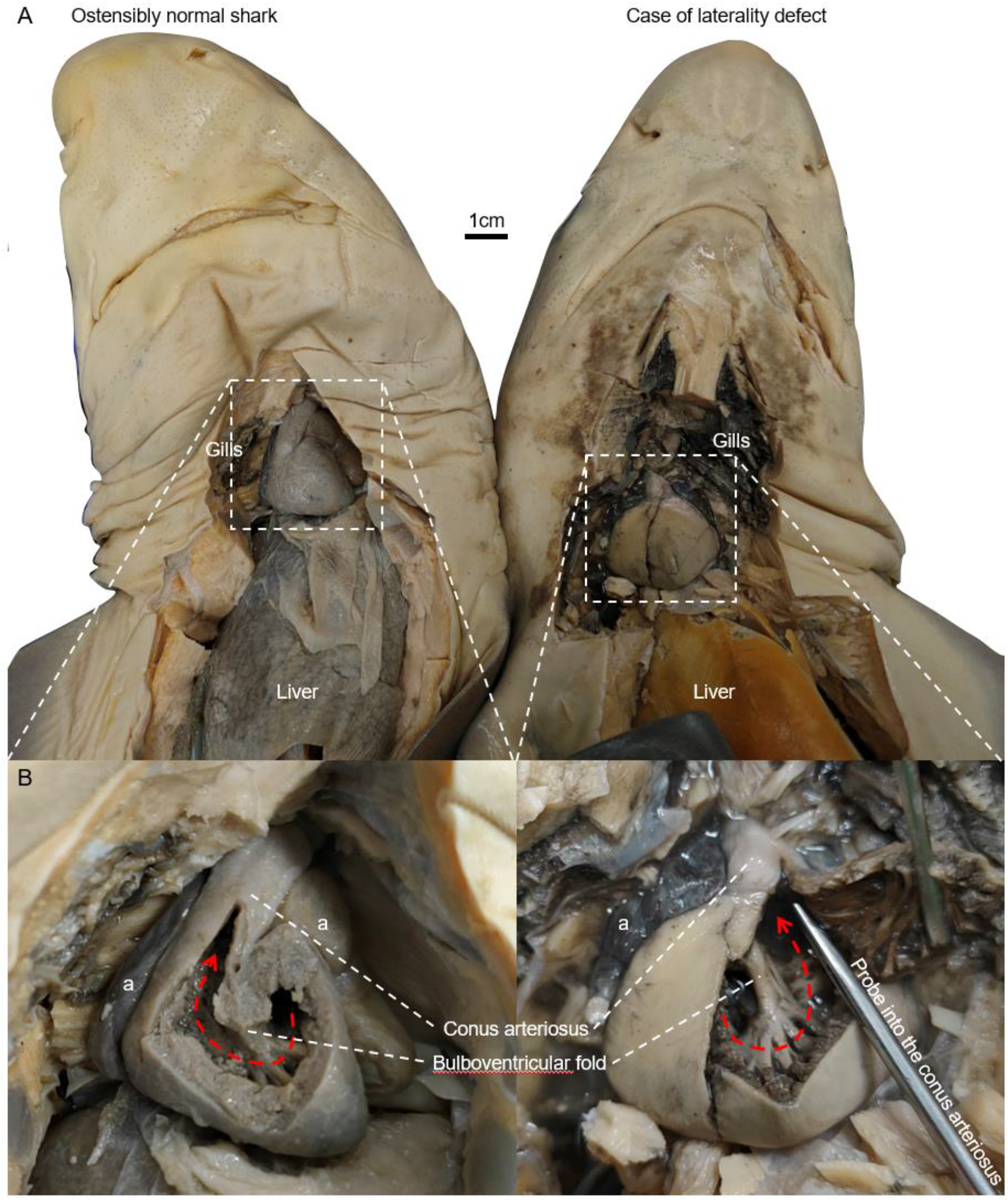
Spiny dogfish with laterality defect. **A**. The shark that exhibited signs of laterality defects (right panel) looks normal. **B**. The heart of the case specimen shows all the structural components expected but the outflow tract is located to the left. Thus, it mirrors the normal setting (compare the dashed red arrows).

The heart of cartilaginous fish has a pronounced left-right asymmetry. The ventricular inlet, or atrioventricular canal, is in the dorsal-left part of the ventricular mass (Sanchez-Quintana and Hurle 1987; Icardo 2017; Hirasaki et al 2018). Blood ejected from the ventricle exits on the right, into the myocardial conus arteriosus (Grimes and Kirby 2009; Lorenzale et al 2018). There is then a pronounced loop in the central cavity of the ventricle, around the so-named bulboventricular fold (Goor et al 1972; Van Mierop and Kutsche 1984; Icardo et al 2004; Jensen et al 2013). This configuration, conserved through evolution, is the retention of the cardiac looping that took place during embryogenesis (Pelster and Bemis 1991). On the luminal side of the ventricle, the bulboventricular fold appears as a large ridge-like aggregate of trabecular muscle (Figure 1B). The bulboventricular fold was readily seen both in the normal hearts and in the heart with the laterality defect (Figure 1B). This suggests that the ventricle of the case specimen contained all the normal structural components. However, whereas the conus arteriosus is normally found to the right of the bulboventricular fold, the conus arteriosus of the case specimen was located to the left of the bulboventricular fold (Figure 1B).

The high-resolution MRI volume rendering recapitulated the anatomy of the whole shark and the inner anatomy of the ventricle (Figure 2A-B). The resulting 3D model of the heart (Figure 2C) revealed a right-sided atrioventricular canal and the left-sided position of the conus arteriosus in the ventricular mass (Figure 2D). This confirmed the mirrored anatomy of the ventricle. The upstream chambers, the atrium and sinus venosus, did not exhibit clear left-right asymmetries, but this is also the case in normally formed hearts. Consequently, the possible existence of lateral deviations could not be determined. The 3D model is available as an interactive pdf (Supplementary Figure 1).

**Figure 2.**
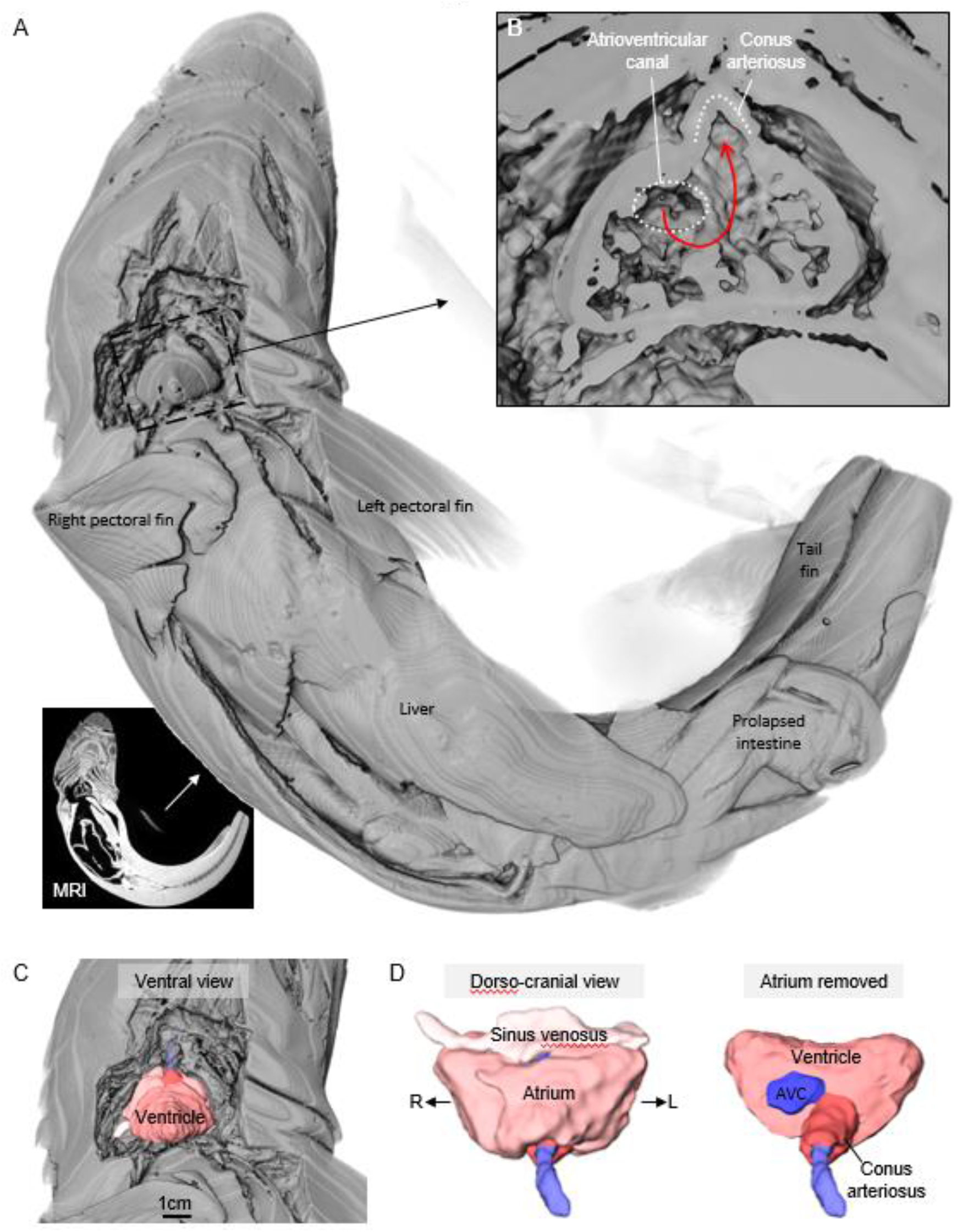
MRI of spiny dogfish with laterality defect. **A**. Volume rendering of the whole shark in ventral view, based on MRI. **B**. The reconstructed ventricle recapitulates the macroscopic inspection shown in Figure 1B. **C**. 3D-model of the heart. Ventral view. **D**. 3D-model of the heart. Cranial view. Removal of the atrium reveals the left-sided position of the atrioventricular canal (AVC). L, left; R, right.

Based on the material of previous studies on the Adriatic sturgeon (Icardo et al 2002, 2004), we report here a cardiac chamber topology defect in a specimen of 90 cm TL (Figure 3A). In addition, out of the approximately three hundred alevins of sturgeon that were previously analyzed to study heart development (Icardo et al., 2004, 2009), about ten per cent exhibited cardiac defects including mirror-looping of the heart. Several examples are shown in Figure 3B. The incidence of inverted heart in those sturgeons is estimated to be between one and two per cent. This incidence is approximately similar to that reported for developing *Xenopus* frogs (van Veenendaal et al 2013), but it is two orders of magnitude more frequent than the approximate 1:10.000 occurrence of dextrocardia reported in humans (Bohun et al 2007). Of note, L-loop hearts were found more frequently in younger than in older alevins. This suggests either the malformation is detrimental in later stages, and, or, it associates with severe extracardiac malformations which, at least in human, can occur (Bohun et al 2007).

**Figure 3.**
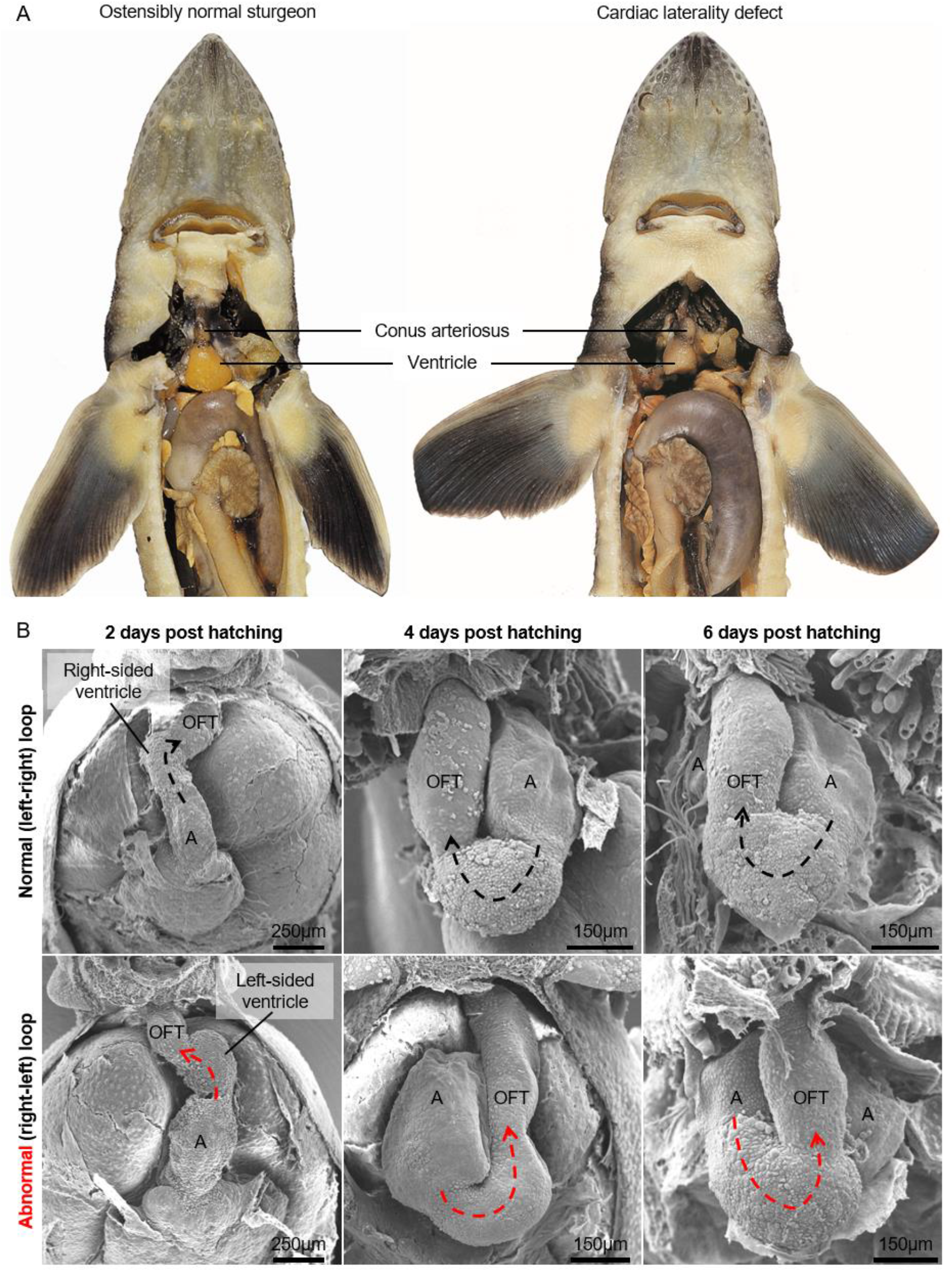
Adriatic sturgeon (*Acipenser naccarii*) with laterality defects. **A**. The adult sturgeon with signs of laterality defects (right panel) looks normal. However, the ventricle is displaced to the right whereas the outflow tract is displaced to the left. **B**. Hearts of sturgeon alevins with normal, R-loop of the ventricle (top row) and cases of mirrored looping (bottom row). A, atrium; OFT, outflow tract.

In conclusion, left-right mirrored hearts occur in both farmed and wild fish. Our findings indicate that laterality defects can easily be identified in cartilaginous fish due to heart asymmetry. These defects may be more difficult to identify in bony (and specially in teleost) fish due to the more left-right symmetric arrangement of the heart chambers.

## Acknowledgements

The authors wish to thank A. Domezain, from the Piscifactoría “Sierra Nevada” at Riofrío, Granada, Spain, for generous access to the sturgeon material. Jaco Hagoort’s help in generating the 3D model is much appreciated.

