## Supplementary Figure 1 for "Laterality defect of the heart in non-teleost fish"

### Information on the use of this interactive 3D-PDF

All structures and cavities
Sinus venosus
Atrium
Ventricle
Conus arteriosus
Sinuatrial junction
Lumen of atrium
Atrioventricular canal
Lumen of ventricle
Lumen of outflow tract

Selected structure:

#### Selection of structures

The top left panel contains buttons to show or hide (groups of) structures, or to make them transparent.

**button example:** show transparent hide Atrium

After a single click on a 3D structure, the structure will be highlighted and the name of the structure will appear below "Selected structure". With the buttons next to the structure name, the appearance of this structure can be changed. Clicking next to the 3D object will deselect the structure.
For more advanced selection options, right-click on the 3D model and choose: “Show Model Tree”.

#### The intended use of this 3D model

This model of the heart is based on few pixels of original data compared to the number of structures and cavities that are reconstructed. Accordingly, the model is somewhat crude. For example, the sinus venosus is represented as a solid object, rather than a cavity surround by a thin wall, and the atrial wall is rendered much thicker than it actually is so that it surrounds the atrial cavity without gaps in the wall. Given short-comings such as these, the main purpose of the model is to give an impression of the position of the major structures and cavities, in particular the right-dorsal position of the atrioventricular canal and the left-cranial emergence of the conus arteriosus from the ventricular mass.

#### Technical Notes

View this PDF file in a recent version of Adobe Acrobat Reader: https://get.adobe.com/reader/

3D interaction is only possible on MS Windows or Mac OS. Javascript and playing of 3D content must be enabled.

*Edit, Preferences* to ensure the following:

1) In *JavaScript*
   - enable *Enable Acrobat JavaScript*
2) In *Multimedia &3D*
   - enable *Enable playing of 3D content*
   - disable *Show 3D Orientation Axis*
   - *Optimization Scheme for Low Framerate: None*

Amsterdam UMC
University Medical Centers

Department of Medical Biology

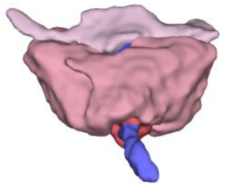

Dorso-cranial view

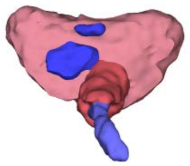

View of atrioventricular canal

### Left-right mirrored shark heart

All structures and cavities

Sinus venosus

Atrium

Ventricle

Conus arteriosus

Sinuatrial junction

Lumen of atrium

Atrioventricular canal

Lumen of ventricle

Lumen of outflow tract

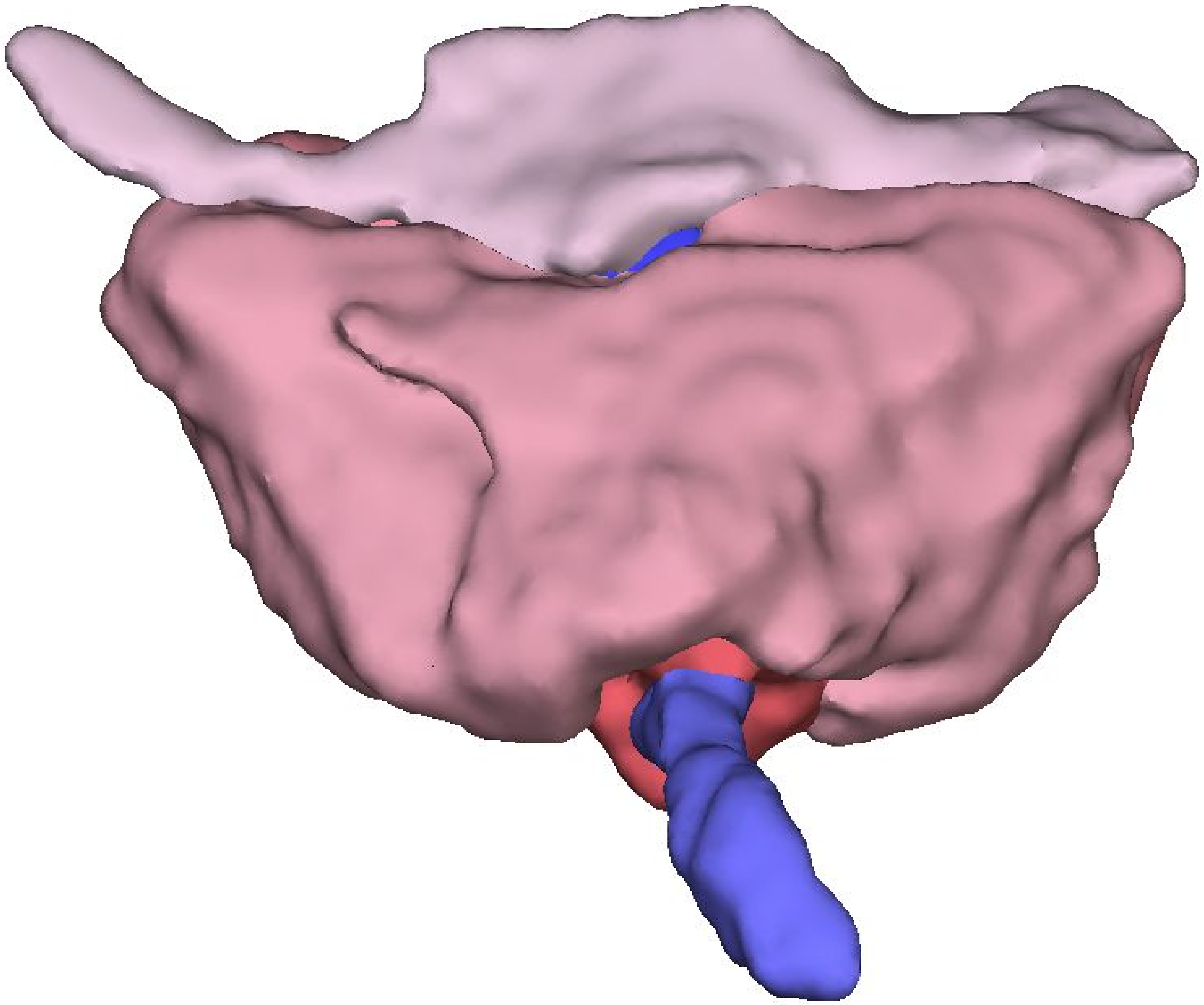

Selected structure:

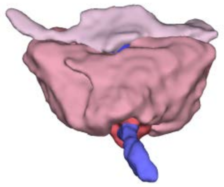

Dorso-cranial view

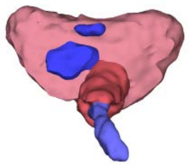

View of atrioventricular canal

Amsterdam UMC
University Medical Centers

Department of Medical Biology
